# Invadopodia-Mediated Remodeling of the Lymphatic Endothelium Drives Cancer Cell Lymphatic Dissemination and is Regulated by a CCR7-Vav2-Rac3 Signaling axis

**DOI:** 10.1101/2025.10.15.682688

**Authors:** Sumreen Javed, Merlyn Emmanuel, Xuewei Chen, Kendal Ruzicki, Esther Afolalan, Nazarine Fernandes, Sepideh Soukhtehzari, Brent DG Page, Karla C. Williams

**Author notes:** Equal Contribution (Sumreen Javed, Merlyn Emmanuel).

## Abstract

Cancer cell invasion across a lymphatic endothelial barrier and subsequent colonization of regional lymph nodes often marks the first stage of metastatic dissemination. Lymphatic vessels facilitate the escape of cancer cells and can support further metastatic dissemination. Although the clinical and experimental evidence supports a role for lymph node metastases in promoting metastatic spread, the mechanism employed by cancer cells to navigate a lymphatic endothelium and enter lymphatic vessels is poorly characterized. To investigate this, we assessed the interactions between cancer cells and lymphatic endothelial cells and found that Tks5-positive structures, termed invadopodia, remodel lymphatic endothelial junctions. Loss of Tks5 impaired cancer cell invasion across a lymphatic endothelium and significantly reduced lymph node and lung metastasis in a mouse model of breast cancer progression. Surgical removal of the axillary and brachial lymph nodes prior to orthotopic cancer cell injection resulted in a significant reduction in lung tumor burden, further demonstrating the significance of lymph node metastases to metastatic tumor burden. Next, using breast cancer patient primary tumors we found that elevated expression of CCR7, a chemokine receptor, significantly associated with lymph node metastasis. CCR7 localized to invadopodia and promoted cancer cell invasion across lymphatic endothelium, both in the presence and absence of its canonical ligand CCL19. Tyrosine phosphorylation of CCR7 directed the recruitment of Vav2 and activation of Rac3. Our findings highlight a role for lymphatic metastases in promoting distant metastasis and establish a mechanism by which cancer cells breach a lymphatic endothelium.

## INTRODUCTION

Metastasis, the dissemination of cancer cells from the primary tumor, is the main cause of cancer related death. The process of cancer cell dissemination is a complex, multistep, process involving local cancer cell invasion at the primary tumor site, intravasation, into, and extravasation, out of, the hematogenous or lymphatic system, and colonization at secondary sites.^1-4^ Key to this process is the ability of cancer cells to escape the primary tumor and enter circulatory systems, which transits them throughout the body. Both the hematogenous system and lymphatic system provide routes of cancer cell dissemination.^5^ While dissemination through the hematogenous system has long been considered the primary route of metastatic spread, increasing evidence suggests that the lymphatic system actively contributes to this process.^6-11^

Lymph nodes are often the first site of metastasis and are critical for tumor staging and prognosis in multiple cancers. Correlative evidence suggests a link between lymph node metastasis and peripheral metastasis,^12^ and studies in mice have demonstrated that lymph node metastases actively contribute to the metastatic seeding of distant organs.^7,9^ Intriguingly, a clinical study investigating evolutionary trajectories of metastatic spread delineated two distinct routes of metastasis: primary tumor (hematogenous) and regional lymph node metastases (lymphatic).^6^ This confirms that the lymphatic system can serve as an intermediate step for distant dissemination. Critically, these works describe distinct prognostic and predictive potential related to the route of metastasis as characterized by poor patient outcomes and reduced response to immunotherapy in the lymphatic route. Taken together, these studies highlight that the lymphatic-mediated dissemination pathway is biologically relevant. To expand on this work, additional studies are needed to explore the mechanism of cancer cell invasion into the lymphatic system.

Cancer cells navigate endothelial barriers to enter into circulatory systems. This process has been well-described for hematogenous dissemination and is regulated by subcellular structures termed invadopodia.^13-16^ Invadopodia are dynamic actin-rich sub-cellular protrusions classically described based on their ability to remodel and degrade the extracellular matrix (ECM). It has also been demonstrated that invadopodia can apply a physical force to their surroundings and can interact with endothelial cell junctions to penetrate across a vascular endothelium.^13^ Invadopodia have been shown to respond to chemotactic ligands by increasing cancer cell extravasation rates into ligand rich microenvironments.^17^ Whether invadopodia support cancer cell navigation through a lymphatic barrier is undefined.

Here, we evaluate the mechanism by which cancer cells breach a lymphatic endothelium, facilitating dissemination into the lymphatic system, and exploit chemotactic cues to promote this process. We examine the interaction of breast cancer cells with a lymphatic endothelium and, through loss of invadopodium regulatory protein, Tks5 (tyrosine kinase family with five Src homology 3 (SH3) domains), we evaluate lymphatic endothelial junction remodeling and invasion. We assess the role of Tks5 in metastatic dissemination and the contribution of lymph node metastases to lung tumor burden. We identify a correlation between CCR7 expression and increased metastatic lymph node tumor burden. Mechanistically, we examine the role of the CCR7 in facilitating invadopodia-based invasion of the lymphatic endothelium and delineated the downstream CCR7 signaling pathway that promotes cancer cell dissemination.

## RESULTS

### Invadopodia Interact with and Modify Lymphatic Endothelial Junctions to Mediate Invasion Across a Lymphatic Endothelium

To investigate the role of invadopodia in cancer cell dissemination through the lymphatics system we generated an in vitro, primary, lymphatics model using human dermal lymphatic cells (HDLEC). Generation of a primary lymphatic endothelium was validated by immunostaining of tight junction marker, ZO-1, and adherens junction marker, VE-Cadherin (Fig. 1A and B). To evaluate the interactions of cancer cells with a lymphatic endothelium, in the context of invadopodia, we generated Tks5-GFP stables using the breast cancer cell line MDA-MB-231. Tks5 was selected as an invadopodia marker due to its established, exclusive, localization at invadopodia and essential function as a core adaptor protein required for invadopodia maturation and stabilization.^18-20^ MDA-MB-231-Tks5-GFP cells were co-cultured on the lymphatic endothelium (Fig. 1C) and their interactions were observed over time. Invadopodia-rich punctae readily formed by one hour and were observed to interact with the lymphatic junctions (Fig 1.D). Lymphatic cells appear to extend projections around the cancer cell as shown by VE-Cadherin membrane staining wrapping around the cancer cell (Fig.1D and E). The localization of Tks5-GFP punctate at sites of lymphatic endothelial junctions demonstrates that invadopodia interact with lymphatic cell junctions and therefore they may mediate a remodeling of lymphatic cell adhesions in support of transendothelial migration through a lymphatic endothelial barrier. To further evaluate the role of invadopodia in mediating cancer cell-lymphatic endothelial cell interactions, we generated control and stable Tks5 knockdown (KD) cell lines (Fig. 1F). As expected, loss of invadopodia formation was found for Tks5-KD cells using a standard invadopodia formation assay (Supplemental Fig. 1A and B). Live cell imaging was performed 30min post-addition of cancer cells to the lymphatic endothelium (T=0) (Fig. 1G, Supplemental Movie 1 and 2). Control cells were observed to migrate on lymphatic endothelial junctions (Fig. 1G, top row: i-iv), arrest (Fig. 1G, top row: iv), and modify junctions to invade into the endothelium (Fig. 1G, top row: v-vi). Tks5-KD cells did not exhibit any impairment in migration and migrated along endothelial junctions similar to control cells (Fig. 1G, bottom row: i-iv). However, Tks5-KD cells appeared to exhibit defects in modifying endothelial junctions and did not penetrate the lymphatic endothelial barrier (Fig. 1G, bottom row: v-vi).

**Figure 1:**
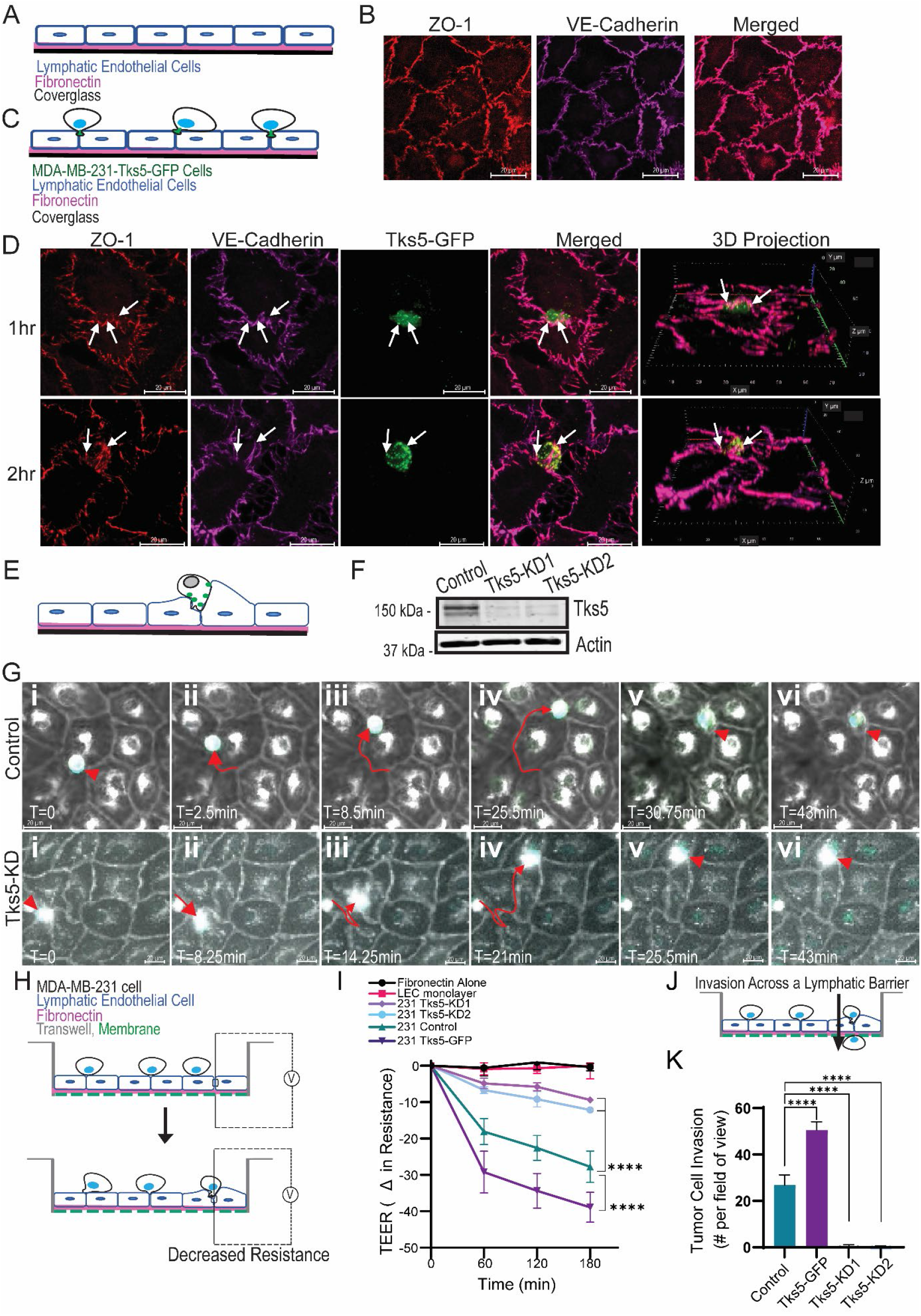
Invadopodia interact with and remodel lymphatic endothelial junctions to invade across a lymphatic endothelium. **(A)** An animated diagram detailing the set-up of a human lymphatic endothelial cell monolayer system. **(B)** A representative image of the lymphatic monolayer stained for tight junction protein ZO-1 and adherens junction protein VE-Cadherin. **(C)** An animated diagram detailing the set-up of the lymphatic endothelial monolayer system incubated with cancer cells. **(D)** Representative images of a Tks5-GFP stable MDA-MB-231 cell incubated on the lymphatic monolayer for 1 hour and 2 hours. Arrows point to Tks5-GFP punctae colocalized with lymphatic tight junction and adherens markers ZO-1 and VE-Cadherin. **(E)** Animated diagram illustrating invadopodium (Tks5-GFP;green punctae) interacting with a primary lymphatic endothelium. **(F)** MDA-MB-231 cells were used to generate Tks5 knockdown cell lines. Representative western blot of Tks5 expression are shown. **(G)** Images from live cell experiments where GFP-MDA-MB-231 control and Tks5 knockdown cell lines added to a lymphatic endothelium and 30-45min post-addition, imaged for 1hr. Lymphatic cell membranes were stained with Cell Mask Orange prior to the addition of MDA-MB-231 cells (light grey; outlining cells). MDA-MB-231 nuclei were stained with Hoechst prior to incubation. Arrowhead at time zero point to an MDA-MB-231 cell (white/blue structure). Arrows point to the movement of cells across the endothelium. Arrowhead at later timepoints indicate arrested cell. Arrowhead pointing to control cells show changes to the lymphatic endothelial junctions/membrane (diffuse light-grey staining). **(H)** Animated diagram depicting cancer cell modification of lymphatic endothelium junctions and the corresponding change to endothelial resistance. **(I)** Transendothelial electrical resistance (TEER) measurements over time for a lymphatic endothelium when incubated with control, Tks5 knockdown, or TKS5-GFP cell lines. Two-way ANOVA. **(J)** Animated diagram depicting cancer cell invasion across a lymphatic endothelium. **(K)** Quantification of MDA-MB-231 control, Tks5 knockdown, or Tks5-GFP cell line invasion across a lymphatic endothelium. One-way ANOVA. N=3. Scale bar= 20µm. Mean ± SEM. Asterisk denotes significance.

To quantify these observed changes, we measured the transendothelial electrical resistance (TEER) of the lymphatic endothelium in the presence of control and Tks5-KD cancer cells. TEER is a non-invasive, established, method to measure the integrity of tight junctions across an endothelium (Fig. 1H). Control or Tks5-KD cells were co-culture on a lymphatic endothelium and modifications to lymphatic endothelial junctions was measured (Fig. 1I). Control cells readily modified the tight junction of the lymphatic endothelium, whereas this was lost in the Tks5-KD cell lines. Next, we evaluated cancer cell invasion through the lymphatic barrier (Fig. 1J,K). The ability of Tks5-KD cell lines to traverse across a lymphatic endothelium was significantly impaired compared to control (Fig. 1K). To further highlight the role of Tks5 in promoting lymphatic endothelial remodeling and invasion, we overexpressed Tks5 in the MDA-MD-231 cell line. A significant increase in the rate of tight junction remodeling and invasion were found when Tks5 expression was increased (Fig. 1I and K). We also inverted our lymphatic endothelial barrier model and tested the ability of control and Tks5-KD cell lines to modify lymphatic endothelial junctions. Tks5-KD demonstrated a significant reduction in the ability to remodel lymphatic endothelial junctions (Supplemental Fig. 2A and B). To determine whether the reduction in endothelial junction remodeling and invasion across the lymphatic endothelium was a result of impaired cell migration, control and Tks5-KD cell line cell migration was assessed. No significant difference in migration was found across the cell lines Supplemental Fig. 3A-D). Next, using two addition breast cancer cell lines (hormone positive MCF7 cell line and HER2-positive 21MT-1 cell line), we evaluated the role of Tks5 in mediating lymphatic endothelial junction remodeling and invasion across a lymphatic endothelium. Loss of Tks5 impaired cancer cell lymphatic endothelial junction remodeling and invasion across a lymphatic endothelium in both cell lines (Supplemental Fig. 4A-F).

### Tks5 Expression Drives Lymph Node Metastasis and Lymph Node Metastases Increase Tumor Burden in the Lung

In order to evaluate the role of invadopodia in lymph node metastasis in vivo, we analyzed Tks5 expression in primary breast tumors and matched lymph node metastases from 47 individuals diagnosed with invasive ductal carcinoma. Metastatic breast cancer cells in the lymph node were found to have a significant increase in Tks5 expression relative to their matched primary tumor (Fig. 2A and B). Next, we evaluated a breast cancer cell line derived from a lymph node metastases: the MDA-MB-231LN cell line. The MDA-MB-231LN cell line was derived from the parental MDA-MB-231 cell line, stably expresses firefly luciferase, and demonstrates homing to the lymph nodes.^21^ An evaluation of MDA-MB-231LN Tks5 expression and invadopodia formation identified significantly higher expression of Tks5 and higher rates of invadopodia formation in the lymph node homing cell line relative to the parental cell line (Fig. 2C-F). MDA-MB-231LN cells were also able to modify lymphatic endothelial junctions at a faster rate relative to MDA-MB-231 cells (Fig. 2G) and demonstrated higher rates of invasion across a lymphatic endothelium (Fig. 2H). To determine if the lymph node homing cell line utilized invadopodia to navigate a lymphatic endothelial barrier, control and Tks5 knockdown (KD) cell lines were generated (Fig. 2I). Cell lines were evaluated for their ability to modify lymphatic endothelial junctions and traverse a lymphatic endothelial barrier. Loss of Tks5 significantly impaired the ability of the lymph node homing breast cancer cells to modify lymphatic endothelial junctions and invade across a lymphatic endothelial barrier (Fig.2J and K).

**Figure 2:**
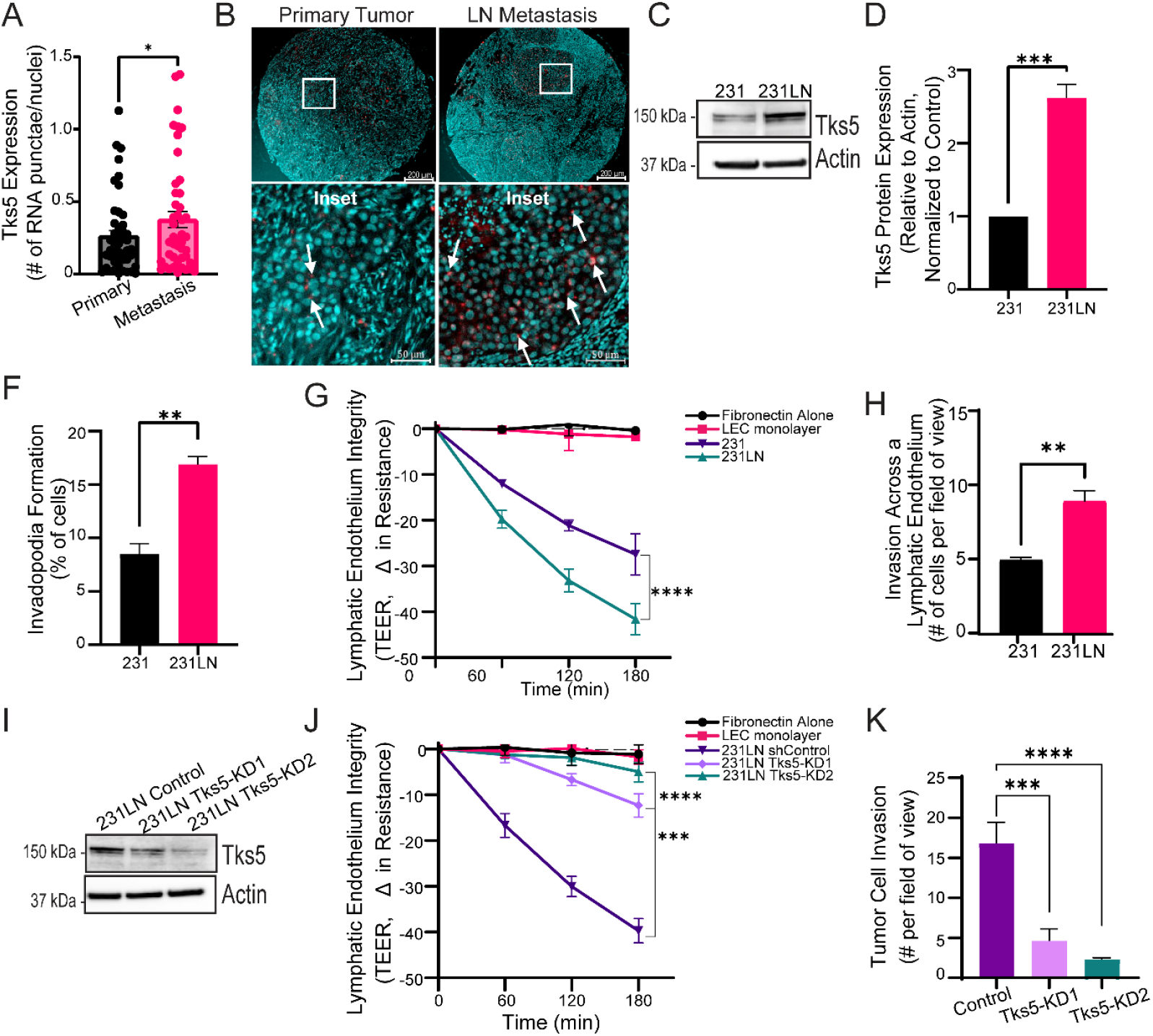
Tks5 expression in elevated in patient lymph node metastases and a metastatic lymph node-derived breast cancer cell line aggressively remodels lymphatic junctions. **(A-B)** Breast tissue microarray containing tissue cores from primary tumors and matched lymph node metastases were stained for Tks5 mRNA. **(A)** Quantification of the number of Tks5 RNA-punctae per nuclei in patient primary tumor (Primary) and breast cancer lymph node (LN) metastases (Metastasis). Paired t-test. **(B)** Representative image of Tks5 expression (red punctae) in primary tumor and matched LN metastasis (cell nuclei are shown as cyan). Arrows point to Tks5 positive punctae. Scale bar = 200 µm (cores) and 50 µm (insets). **(C)** Representative western blot of Tks5 expression in MDA-MB-231 cell line and LN metastatic derivative cell line MDA-MB-231LN. **(D)** Quantification of western blots as shown in **(C)**. N=3. One Sample t-test. **(F)** Cell lines were plated on gelatin-coated coverslips for 4 h, fixed, permeabilized, and stained with Alexa488-phalloidin to stain F-actin. Cells were image using a fluorescent microscope. Cells overlaying spots of gelatin degradation, co-localized with F-actin, were classified as positive for invadopodia. The percentage of cells forming invadopodia is shown; 20 independent spots per sample were counted. N=3. Student t-test. **(G)** Transendothelial electrical resistance (TEER) measurements over time for a lymphatic endothelium when incubated with MDA-MB-231 or MDA-MB-231LN cell lines. N=3. Two-way Anova. **(H)** Quantification of MDA-MB-231 or MDA-MB-231LN cell line invasion across a lymphatic endothelium. One-way Anova. **(I)** MDA-MB-231LN cell line was used to generate cell lines stably expressing a constructs targeting Tks5 (231LN Tks5-knockdown 1 [KD1] and 2 [KD2]) or control, non-targeting construct (231LN Control). Representative western blot is shown. **(J)** Transendothelial electrical resistance (TEER) measurements over time for a lymphatic endothelium when incubated with MDA-MB-231LN control and Tks5 knockdown cell lines. Two-way Anova. **(K)** Quantification of MDA-MB-231LN control and Tks5 knockdown cell lines invasion across a lymphatic endothelium. One-way Anova. Mean ± SEM. N=3. Asterisk denotes significance.

To assess the metastatic capacity of MDA-MB-231LN control and Tks5-KD cell lines, cell lines were orthotopically injected into the 2^nd^ mammary fat pad of female NOD SCID gamma (NSG) mice and tumor growth was monitored by bioluminescence imaging (BLI) (Fig.3A). Tks5-KD tumors exhibited a slightly slower growth rate which was significantly different on day 7 (Fig. 3B). Since we wanted to measure metastatic tumor burden in the lymph node and lungs while mitigating the effects of tumor growth, Tks5-KD mice were monitored until their primary tumor mass matched that of control tumors. At endpoint, no significant difference was found between control and Tks5-KD primary tumors (Fig. 3A and B).

**Figure 3:**
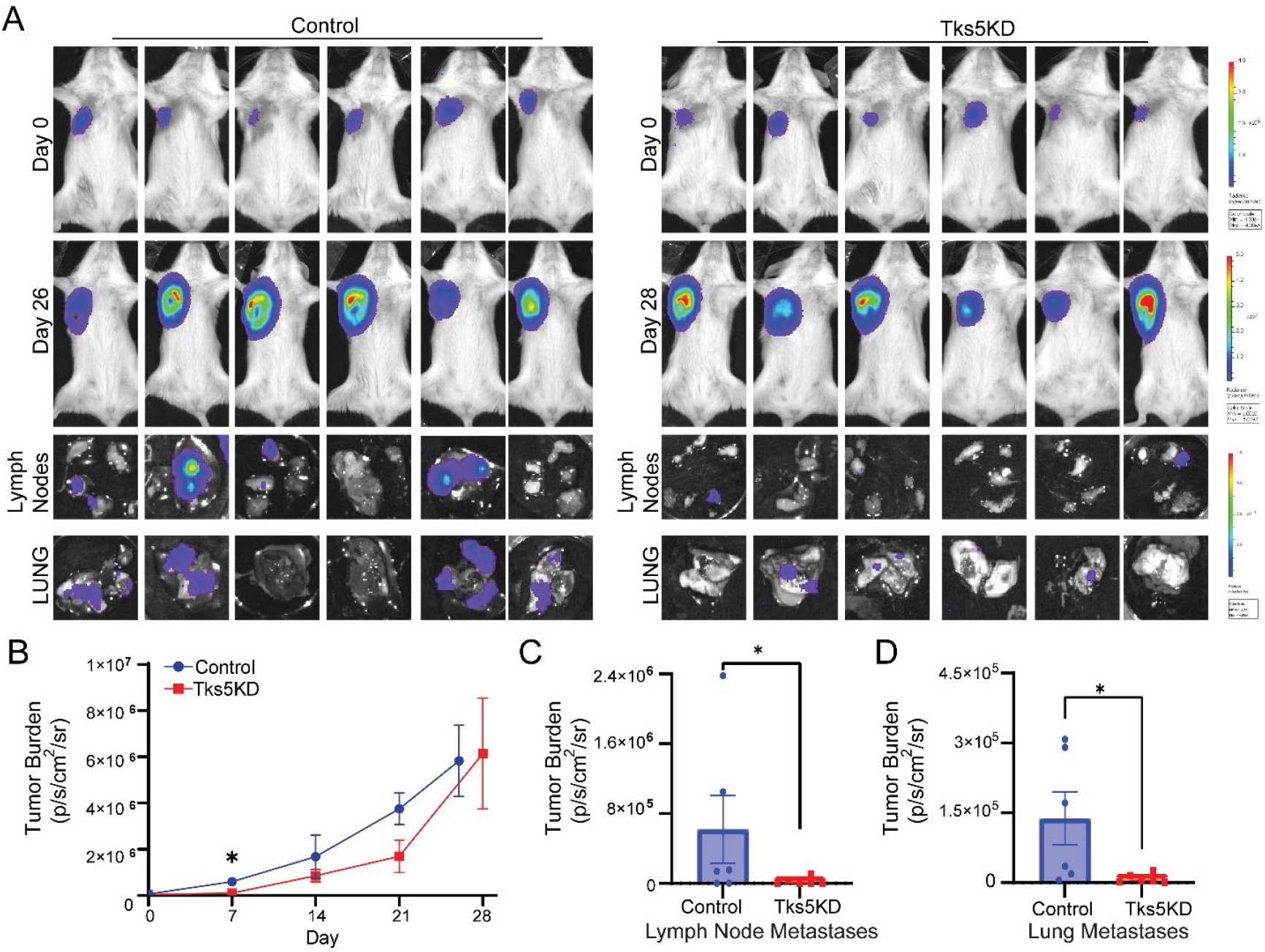
Tks5 mediates tumor cell dissemination into the lymphatic system and the lungs. **(A)** MDA-MB-231LN control and Tks5 knockdown cell lines were mammary fat pat (m.f.p.) injected into NSG mice. Mice were i.p. injected with luciferin and bioluminescence imaging (BLI) was performed to ensure equivalent cell injection numbers at experimental start point and to monitor control and Tks5 knockdown cell line tumor growth overtime. Images from BLI of mouse mammary tumors post-m.f.p. injection (top row; n=6 control and n=6 Tks5 knockdown) and at endpoint for control (Day 26) and Tks5 knockdown (Day 28). Bottom two rows: images from end-point ex vivo BLI of lymph nodes and lungs at study endpoint. **(B)** Quantification of bioluminescence signal from control and Tks5 knockdown primary tumors overtime to monitor tumor growth. One-way ANOVA. Metastatic tumor burden, as measured by BLI, in the **(C)** lymph nodes and **(D)** lungs from mice injected with control or Tks5 knockdown cell lines at study endpoint. Means ± SEM, asterisk denotes significance, Student’s T-test.

The right axillary and brachial lymph nodes, proximal to the tumor, and the lungs were removed for ex vivo imaging (Fig. 3A, bottom two rows). A significant reduction in metastatic tumor burden in the lymph node and lungs was found for mice with Tks5-KD tumors compared to control tumors (Fig. 3C and D). Overall, this demonstrates that Tks5 mediates metastasis to the lymph nodes and lungs, strongly suggesting that this process is regulated by invadopodia.

Finally, we wanted to assess the contribution of lymph node metastases to the overall lung tumor burden. As we found that MDA-MB-231LN cells invaded and colonized the brachial and axillary lymph nodes, proximal to the 2^nd^ mammary fat pad, of NSG mice we surgically removed these lymph nodes prior to the injection of cancer cells. Surgery, or sham surgery, was performed and mice were given 4 days to recover prior to injection of MDA-MB-231LN cells into the 2^nd^ mammary fat pad. Tumor growth was monitored till endpoint (Day 23) and no significant difference in tumor growth was found between the sham surgery group or lymph node removal surgery group (Fig. 4A and B). We noted that the primary tumor burden for the surgery groups increased at a faster rate compared to that of the control/Tks5-KD study. The tumor burden of mice that underwent surgery reached a total radiance of ∼4.9x10^6^ by day 14 (Fig. 4C) compared to that of control mice at ∼1.6x10^6^ on day 14 (Fig. 3B); due to primary tumor burden, the surgery study endpoint was set at day 23. Lungs were removed at endpoint and tumor burden was assessed by by ex vivo BLI (Fig. 4A and B). A significant reduction in metastatic tumor burden was identified for the lymph node removal surgery group compared to the sham surgery group (Fig. 4D). This suggests that lymph node metastases promote metastatic tumor burden in the lung.

**Figure 4:**
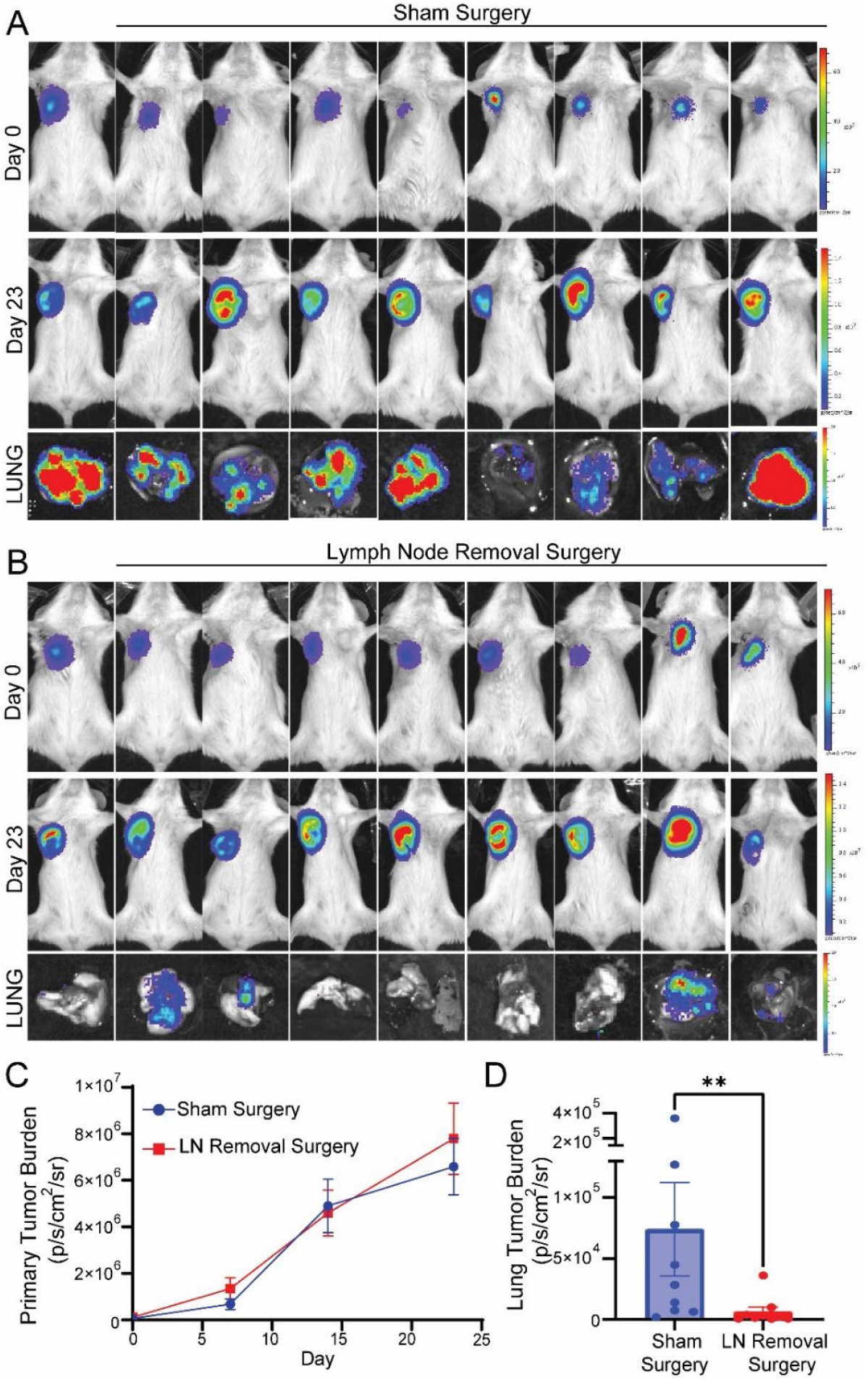
Tumor cell dissemination into the lymphatic system promotes lung tumor burden. NSG mice underwent either: **(A)** mock surgery (sham) or **(B)** had surgical removal of their brachial and axillary lymph nodes (lymph node removal). **(A and B)** Post-surgery, MDA-MB-231LN cells were mammary fat pad injected into mice and BLI was performed. Images of bioluminescence signal from the primary tumor on Day 0 and Day 23 are shown. Bottom row shows images of bioluminescence signal from the lungs, ex vivo BLI, of sham and lymph node removal groups at endpoint. **(C)** Bioluminescence signal measured overtime to monitor tumor growth. **(D)** Metastatic lung tumor burden, as measured by BLI, in the lungs of sham surgery and lymph node removal surgery mice. Mean± SEM. Asterisk denotes significance. Student’s T-test.

### CCR7 Promotes Invadopodia Formation and Cancer Cell Invasion Across a Lymphatic Endothelium

Cancer cells can exploit chemotactic cues present in the microenvironment to promote dissemination and colonization of specific niches. Our prior work demonstrated that invadopodia are chemotactic structures capable of guiding cancer cell invasion into distinct microenvironments.^17^ Given the well-characterized role of CCR7 in dendritic cell lymph node homing,^22,23^ we investigated a potential role for CCR7 in promoting invadopodia-dependent cancer cell invasion into the lymphatic system. CCR7 expression was evaluated in breast cancer cell lines and primary breast tumor tissue. The lymph node homing cell line (MDA-MB-231LN) was found to express significantly higher levels of CCR7 compared to the parental MDA-MB-231 cell line (Fig.5A and B). Next, we evaluated CCR7 expression in primary breast tumors (n=75) (Fig. 5C). A correlation matrix evaluating CCR7 expression in relation to patient clinical characteristics identified the strongest positive correlation with lymph node positivity (Supp. Fig. 5A). CCR7 expression was found to be significantly higher in tumors from patients who had multiple lymph node metastases (Fig. 5D). To further expand on this using a larger cohort, we accessed mRNA expression data from Molecular Taxonomy of Breast Cancer International Consortium (METABRIC).^24,25^ CCR7 primary tumor expression was grouped based on the number of lymph nodes found to be positive for metastases (Fig. 5E). As the number of lymph nodes positive for metastases increased, so did the expression of CCR7 and patients with multiple positive lymph nodes (n=4-9, and 10+) had significantly higher expression of CCR7.

**Figure 5:**
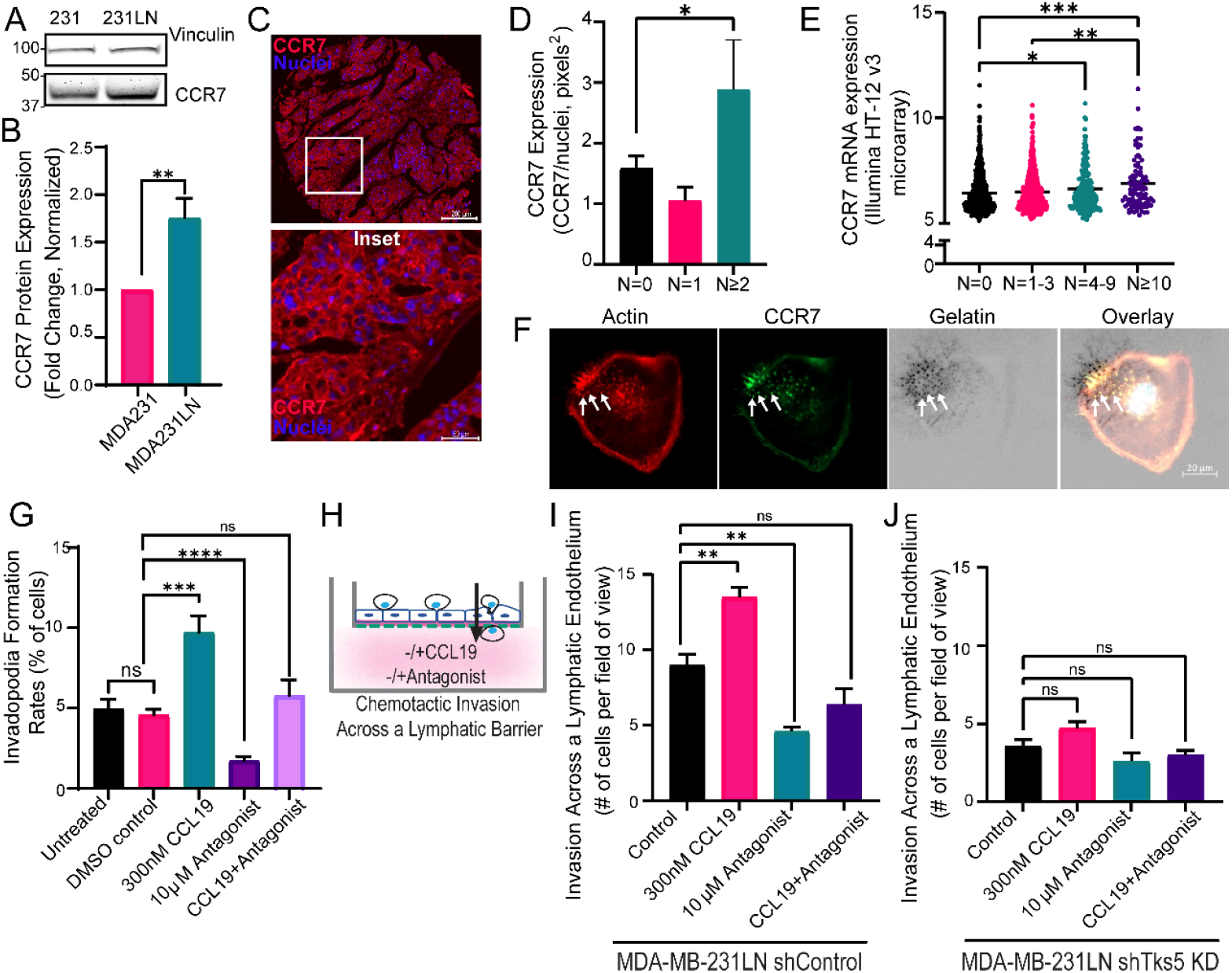
CCR7 expression correlates with lymph node metastases and promotes invadopodia formation and lymphatic endothelium invasion. **(A, B)** CCR7 expression in MDA-MB-231 and MDA-MB-231LN cell lines. Representative western blot of CCR7 expression levels **(A)** used for quantification of CCR7 expression **(B)**. N=3. One Sample t-test. **(C)** Breast tumor tissue microarray immunostained for CCR7. Representative image of CCR7 localization and expression on breast tumor cells positive for CCR7 (red). Nuclei are shown in blue. Scale bar = 200 µm (cores) and 50 µm (insets). **(D)** Analysis of CCR7 expression on tumor cells categorized based on the number of nodes positive for metastatic breast cancer cells. One-way ANOVA. **(E)** METABRIC database extraction of CCR7 mRNA levels from clinical breast cancer samples (n=1866) grouped by number of lymph nodes positive for breast cancer cells. Kruskal-Wallis test. **(F)** MDA-MB-231LN cells were seeded on 568-labeled gelatin coated coverglass, incubated for 4h, fixed and immunostained for CCR7 (green). F-actin (647-phalloidin) is shown as red; 568-labeled gelatin as greyscale, and nuclei are blue (seen as white in the overlay). Invadopodia are noted by F-actin punctae overlaying spots of degradation. Arrows point to CCR7 localization at invadopodia. Scale bar = 20 µm. **(G)** MDA-MB-231LN cells were seeded on 568-labeled gelatin coated coverglass, incubated with control (DMSO), CCL19 ligand, CCR7 antagonist, or CCL19 ligand plus CCR7 antagonist. 3h post-seeding, cells were fixed, permeabilized, and stained for F-actin (488-phalloidin). The percentage of cells forming invadopodia was quantified for each condition. One-way ANOVA. **(H)** Animated diagram depicting breast cancer cell invasion across a lymphatic endothelium as regulated by chemotactic ligand receptor activity. **(I)** Quantification of MDA-MB-231LN cell line invasion across a lymphatic endothelium in the presence, or absence, of CCR7 ligand (CCL19) or CCR7 antagonist. Invasion rates across a lymphatic endothelium for **(J)** MDA-MB-231LN control and **(K)** MDA-MB-231LN Tks5 knockdown cell lines with the addition, or absence, of CCR7 ligand (CCL19) or CCR7 antagonist. N=3. One-way Anova. Mean ± SEM. Asterisk denotes significance.

To evaluate a role for CCR7 in promoting invadopodia-dependent cancer cell invasion across a lymphatic endothelium, CCR7 localization at invadopodia was assessed. CCR7 was readily found to localize to invadopodia (Fig. 5F; white arrows indicate active invadopodia) and stimulation with CCL19 increased the rates of invadopodia formation (Fig. 5G). There was no significant difference in invadopodia formation when cells were stimulated with CCL21 suggesting that CCL19 is the primary ligand functioning through CCR7 to promote invadopodia formation (Supplemental Fig. 6A). To block CCR7 activation we utilized a small molecule allosteric inhibitor of CCR7 (Cmp2105) which binds to the intracellular domain of CCR7 blocking G protein and β-arresting binding. Cancer cells treated with CCL19 in the presence of the CCR7 antagonist did not increase their rates of invadopodia formation (Fig. 5G). In the absence of exogenously added CCL19, CCR7 antagonist dramatically reduced invadopodia formation rates compared to control (Fig. 5F). As CCR7 was found to exhibit activity in the absence of exogenous CCL19, we considered the possibility that breast cancer cell lines produce endogenous CCL19. Cell lysates and conditioned media were assessed for CCL19 expression. CCL19 was not detected in cell lysates or conditioned media (Supplemental Fig. 6B). This demonstrates that CCR7 exhibits activity in the absence of CCL19 and directly contributes to invadopodia formation, or function, which can be amplified with the addition of exogenous CCL19. To evaluate CCR7 function in promoting invadopodia-based cancer cell dissemination across a lymphatic endothelium, control and Tks5-KD cell lines were assessed for chemotactic invasion across a primary lymphatic endothelium (Fig. 5H). Blocking CCR7 activity (Cmp2105) resulted in a significant reduction in cancer cell invasion across the lymphatic endothelium whereas promoting CCR7 activity (+CCL19) significantly increased invasion (Fig. 5I). The Tks5-KD cell line was non-responsive to CCL19 (Fig. 5J). These results strongly suggest that CCR7 functions to promote invadopodia formation in support of cell movement across a lymphatic endothelial barrier (Fig. 5J).

### CCR7 regulates invadopodia function through Rac3-Vav2 duo

CCR7 is a key chemokine receptor known for its central role in dendritic cell migration *via* regulating lamellipodia formation. Our results suggest that CCR7 has a role in regulating cancer cell lymphatic invasion *via* invadopodia. To further validate this, we stably knocked down CCR7 in the MDA-MB-231LN cell line (Fig. 6A) and subsequently measured invadopodia formation rates. Stable knock down of CCR7 led to a marked reduction in invadopodia activity (Fig. 6B). Src kinase is a master director of invadopodia and regulates its formation and activity through the phosphorylation of multiple substrates localized at invadopodia. CCR7 has been shown to be phosphorylated by Src kinase and CCR7.^26,27^ To determine whether CCR7 also undergoes Src kinase mediated phosphorylation in breast cancer cells, we immunoprecipitated CCR7 from MDA-MB-231LN cells and assessed tyrosine phosphorylation using a phosphotyrosine specific antibody. CCR7 was found to be tyrosine phosphorylated (Fig. 6C). Src-mediated phosphorylation of CCR7 creates a docking site for signaling molecules such as Vav1 and SHP2.^26^ Given that phosphorylated CCR7 creates a docking site for signaling molecules, consideration was given for the binding of Vav proteins to CCR7. Co-immunoprecipitation of CCR7 identified a Vav2 interaction. Notably, inhibition of Src kinase, leading to loss of tyrosine phosphorylation of CCR7, diminished Vav2 binding to CCR7, (Fig. 6D) suggesting that the CCR7-Vav2 binding is dependent on Src kinase phosphorylation.

**Figure 6:**
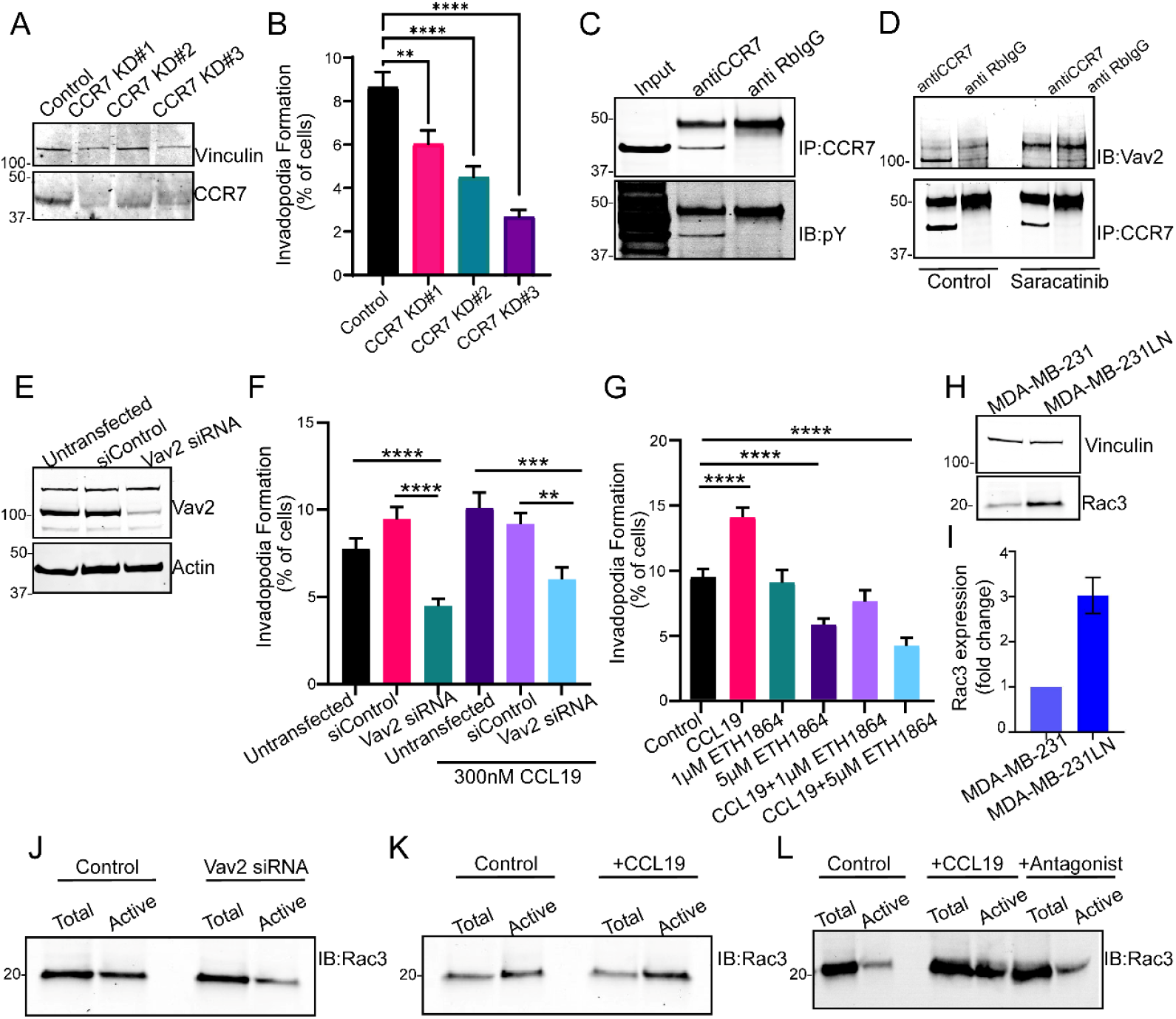
CCR7 regulates invadopodia formation and activity through a Vav2-Rac3 axis. **(A)** The lymph node– homing MDA-MB-231 cells (231LN) was used to generate stable cell lines expressing either a non-targeting control or CCR7-targeting sequence. Western blot of CCR7 expression in control and knockdown cell lines. **(B)** Control and CCR7 knockdown cell lines were seeding on 568-labeled gelatin and rates of invadopodia formation were quantified. N=3. One-way ANOVA. **(C)** CCR7 tyrosine phosphorylation. CCR7 was immunoprecipitated from MDA-MB-231LN cells and analyzed by western blot for tyrosine phosphorylation. **(D)** Src-dependent tyrosine phosphorylation of CCR7 is required for Vav2 binding. MDA-MB-231LN cells were treated with control or Src inhibitor (saracatinib) prior to CCR7 immunoprecipitation. Western blot of co-immunoprecipitated Vav2 and CCR7 are shown. **(E-F)** Transient knockdown of Vav2 reduces invadopodia activity. Control and Vav2 siRNA cells were plated on 568-labeled gelatin and either left untreated or treated with control or CCL19. Rates of invadopodia formation were quantified. One-Way ANOVA. **(G)** Pan Rac inhibitor ETH1864 was added to MDA-MB-231LN cells plated on568-labeled gelatin and rates of invadopodia formation were quantified. One-way ANOVA. **(H**,**I)** Rac3 expression in MDA-MB-231LN cells compared with parental MDA-MB-231 cell. **(H)** Representative western blot of Rac3 expression and **(I)** quantification of Rac3 expression. One Sample T-test. **(J**,**K**,**L)** Rac3 activity in MDA-MB-231LN cells as measured by Pak1-PBD pulldown assays.**(J)** Rac3 activity in control and Vav2 knockdown cells. **(K)** Rac3 activity in the presence of CCL19 or control. **(L)** Rac3 activity in the presence of CCR7 ligand (CCL19), CCR7 antagonist, or control. Mean ± SEM. Asterisk denotes significance.

Given that Vav family of GEFs are critical activators of Rho GTPases that regulate actin cytoskeleton remodeling, transient knockdown of Vav2 was performed (Fig. 6E) and invadopodia activity was evaluated under basal and CCL19-stimulated conditions. Loss of Vav2 markedly impaired invadopodia mediated matrix degradation relative to control cells. Importantly, CCL19 treatment, failed to rescue this defect in Vav2-depleted cells (Fig. 6F). These results indicate that Vav2 is required for CCR7-mediated invadopodia activity. Finally, we assessed the Rac GTPase substrate of Vav2 which can regulate actin cytoskeleton remodeling downstream of CCR7. Initially, we used the small-molecule pan-Rac inhibitor EHT1864 to assess the role of Rac in invadopodia regulation.^28^ Consistent with previous reports,^29,30^ Rac inhibition led to a significant reduction in invadopodia activity, which could not be rescued by CCL19 stimulation (Fig. 6G). These findings confirmed that Rac GTPases are essential for invadopodia function downstream of CCR7. To determine which Rac GTPase mediates this effect, Rac3 expression was assessed and found to be selectively upregulated in the lymph node–homing derivative (MDA-MB-231LN) relative to the parental cell line (Fig. 6H and I). Rac3 activity was assessed using a Pak1-PBD effector pulldown assay to evaluate activity in control and Vav2 knockdown cell lines. Loss of Vav2 lead to a dramatic reduction in active Rac3 suggesting that Vav2 is an activator of Rac3 (Fig. 6J). To confirm that CCR7 activity leads to downstream Rac3 activation, Rac3 activity was assessed in the presence and absence of CCL19. Active, GTP-bound, Rac3 increased upon CCL19 stimulation (Fig. 6K) and this was blocked by CCR7 antagonist Cmp2105 (Fig. 6L). Taken together, these results identify Rac3 as a downstream effector of CCR7 signaling and demonstrate that CCR7 regulates invadopodia function through a Vav2-Rac3 axis.

## DISCUSSION

The findings of this study establish a role for Tks5 in mediating invasion across a lymphatic endothelium and in promoting lymph node metastasis. As Tks5 is essential for the formation and function of invadopodia, our work strongly suggests that invadopodia are essential for cancer cells to navigate and penetrate a lymphatic endothelial barrier. A number of studies have highlighted the role of invadopodia, in vivo, at various stages of cancer cell dissemination. These studies describe an essential role for invadopodia in enabling cancer cells to navigate, and modify, the local microenvironment in support of basement membrane degradation, intravasation and extravasation.^13,31-33^ In addition, invadopodia are chemotactic structures− capable of responding to environmental cues to drive cancer cell invasion into defined microenvironments.^17^ Our study complements and expands this literature by providing supporting evidence for the role of invadopodia in mediating cancer cell dissemination through the lymphatic system. In addition, our work provides evidence for the role for lymph node metastases in promoting metastatic lung tumor burden.

Lymphatic vessels provide key routes that recirculate cells and fluid entering the systemic circulation. Lymphatic vessels also provide an essential route for immune cells to gain access to lymph nodes which plays a critical role in host defense systems.^34^ Lymphatic vessel integrity is maintained through endothelial junctions. VE-Cadherin is a marker of lymphatic endothelial cell junctions and functions to stabilize and maintain endothelial cell-cell adhesion.^35^ Our model was comprised of a primary lymphatic endothelium with continuous VE-cadherin and tight junction, ZO-1, immunoreactivity. Using this model, we have shown a time-dependent interaction of cancer cell invadopodium marker, Tks5, with lymphatic endothelial cell-cell junctions. The Tks5-associated puncta colocalized with with both ZO-1 and VE-Cadherin at the first and second hour. It was also observed that these interactions appear as if the cancer cell is being wrapped around the lymphatics prior to movement through the lymphatic barrier. We attribute the interaction and modification of lymphatic endothelial tight junctions with Tks5-mediated invadopodia activity, as loss of Tks5 impaired endothelial cell-cell junction remodeling across multiple different breast cancer cell lines. A prior report mapping the origin of metastases indicates that lymphatics pre-select an aggressive phenotype of tumor cells and support their dissemination to the secondary site.^6^ Our results provide additional support for this assessment as cancer cell Tks5 expression, and invadopodia formation, are established drivers of aggressive cancer progression.

Cancer cell entry into the lymphatic system could be a passive or active process. Local cancer cell migration could lead to cancer cell interaction with lymphatic vessels and active invasion into collecting vessels. Alternatively, the primary tumor could drive the formation of dysregulated lymphatic capillaries or collecting vessels whereby leaky vessels permit passive cancer cell movement. Further movement out of the lymph node is probable to be an active process whereby cancer cells capable of remodeling lymphatic endothelial tight junctions are able to exit and invade local blood vessels to promote distant metastasis. A number of studies support the role of lymph node blood vessels as a route for systemic dissemination of cancer cells.^7-9^ In addition, a recent study performing single-cell lineage tracing of breast cancer cell dissemination provides further evidence for the lymphatic system as an important route of cancer cell dissemination.^36^ Findings detailed in our study add to this emerging literature by revealing that cancer cells lacking the capacity to remodel lymphatic tight junctions (i.e. loss of Tks5) displayed reduced metastatic spread to the lymph nodes and lungs. In addition, lymph node metastases were found to promote metastatic tumor burden as mice lacking lymph nodes, proximal to the primary tumor, had lower metastatic lung tumor burden. This indicates that lymphatic metastases hold an important role beyond correlative clinical prognostication and actively contribute to, or promote, the formation of distant metastases.

To better understand the process promoting, or supporting, cancer cell invasion across a lymphatic endothelium, we focused on CCR7. CCR7 expression is upregulated on mature dendritic cells and promotes cell migration and homing to lymphatic vessels. Dendritic cells can enter the lymphatics via the capillary vessel, with discontinuous junctions facilitating immune cell transmigration, or collector vessel, with continuous junctions and decreased permeability. Movement into collector vessels promotes rapid migration to lymph nodes and is dependent of CCR7.^37^ Evaluation of CCR7 expression identified elevated expression in a lymph-node homing metastatic breast cancer cell line relative to its parental cell line. In addition, we found elevated expression of CCR7 in primary tumors from breast cancer patients correlate with increased number of lymph nodes positive for metastases. Mechanistically, CCR7 was found to localize to invadopodia. Stimulation with CCR7 ligand CCL19 increased the rates of invadopodia formation. Conversely, a CCR7 antagonist (Cmp2105) dramatically reduced basal invadopodia formation and blocked the increase seen with CCL19 stimulation. The CCR7 antagonist also resulted in a significant reduction in cancer cell invasion across the lymphatic endothelium, while promoting CCR7 activity with CCL19 significantly increased invasion. Critically, we show that this is dependent on invadopodia as loss of Tks5 expression lead to cells being non-responsive to CCL19. These finding strongly suggests that CCR7 functions to promote invadopodia formation in support of cell movement across a lymphatic endothelial barrier.

To delineate the downstream pathway connecting CCR7 activation to invadopodia function we assessed potential CCR7 interacting protein− Vav2. It has been demonstrated that Src tyrosine phosphorylation of CCR7 creates a docking site for adaptor proteins.^26,27^ In mature dendritic cells, Vav1 is recruited to lamellipodia-localized CCR7 leading to Rac1 activation and subsequent cell migration. We considered that Vav proteins are recruited to CCR7 at invadopodia to promote invadopodia formation through Rac-mediated cytoskeleton reorganization. Vav2 recruitment to invadopodia has been demonstrated and was found to be dependent on its SH2 domain.^38^ Vav2 recruitment resulted in Rac3 activation and Rac3 was shown to be enriched at the core of invadopodium early in formation and promoted stability and maturation.^39^ Our analysis of Vav2 and Rac3 identified a CCR7-Vav2 binding event that was dependent on tyrosine phosphorylated CCR7. Loss of Vav2 impaired invadopodia-mediated matrix degradation, and CCL19 treatment failed to rescue this defect. Loss of Vav2 dramatically reduced active Rac3, suggesting Vav2 is an activator of Rac3. Furthermore, active Rac3 increased upon CCL19 stimulation, and this activation was blocked by CCR7 antagonist Cmp2105. Overall, our results describe the localization of CCR7 at invadopodia and detail a novel mechanism whereby CCR7 regulates invadopodia function through a Vav2-Rac3 axis. This pathway clarifies how external chemokine cues are transduced internally to drive the physical, invadopodia-based, activities required for lymphatic invasion.

## Acknowledgements

This work is supported by funding from the Canadian Institutes of Health Research (CIHR) Project Grant awarded to K.C.W. S.S was supported by a CIHR doctoral research award. K.C.W. holds a Tier 2 Canada Research Chair.

## EXPERIMENTAL PROCEEDURES

### Materials and Reagents

All reagents were purchased from either Sigma-Aldrich (St. Louis, MO) or Thermo Fisher Scientific (Nepean, ON, Canada) unless otherwise stated. Antibodies specific for following proteins were purchased from the suppliers indicated: Tks5 (for western blot: Cell Signaling 16619S; 1:250), Tks5 (for Immunofluorescence: Millipore Sigma 09-403), β-actin (for western blot: Sigma A5441; 1:5000), ZO-1 (Thermo Fisher Scientific 33-9100; 1:50), Vinculin (for western blot: Abcam, ab91495, 1:3000), VE-Cadherin (Abcam ab33168; 1:50), CCR7 (for Immunofluorescence (1:500) and Immunoprecipitation (1ug): Novus Biologicals NBP2-67324), CCR7 (for western blot: Abcam ab 191575, 1:500), Vav2 (Proteintech 21924-1-AP,1:500), Rac3 (Invitrogen MA5-53241,1:500), Phosphotyrosine (for western blot: Invitrogen 14-5001-82, 1:1500) and Rabbit Isotype control (Biolegend, Cat. No. 910801). The secondary antibodies (1:3000) and phalloidin labeled with Alexa Fluor 647 (1: 1000) were purchased from Invitrogen (Burlington, Canada). The Alexa 568 Protein Labeling Kit was purchased from Invitrogen (Cat. No. A10238). Lenti ORF particles for Tks5-GFP were ordered from Origene (Rockville Md; RC217487L4V), Tks5 knockout constructs were also purchased from Origene Technologies CAT#: KN417487. Tks5 knockdown plasmids (SH3PXD2A siRNA Lentivector Set, Cat. No. LS143682), CCR7 knockdown plasmids (siRNA Lentivector Set, Cat. No. LS115431) and control plasmid (Cat. No. LV015) were ordered from Applied Biological Materials (Richmond, BC, Canada), and Luciferase-pcDNA3 was a gift from William Kaelin (Addgene plasmid # 18964 ; http://n2t.net/addgene:18964 ; RRID:Addgene_18964 (24). The human Vav2 siRNA SMART pool was purchased from Dharmacon, Horizon Discovery (Cat. No. L-005199-00-0005). The Src Kinase inhibitor, Saracatinib was purchased from Tocris (Cat. No. 7189). Recombinant human CCL19 was purchased from Thermo Fisher Scientific (Cat. No. 300-29B). CCR7 antagonist CCR7 ligand 1 (CCR7-Cmp2105) was purchased from MedChem Express (Cat. No. HY-133073). The transfection reagent, Turbofectin, was ordered from Origene (Cat. No. TF81001) and lipofectamine3000 was purchased from Thermo Fisher Scientific (Cat. No. L3000008). The Pierce Rapid GOLD BCA Protein assay kit was purchased from Thermo Fisher Scientific (Cat. No. A55860). The Protein G Magnetic beads were purchased from NEB (Cat. No. S1430S). The GTP stock was purchased from Thermo Fisher Scientific (100mM, Cat. No. R0461). Magnetic Glutathione Agarose beads were purchased from Thermo Fisher Scientific (Cat. No. 78601). ETH1864 was purchased from Cayman Chemical (Cat. No.17258). The GST PAK1-PBD plasmid was a generous gift from Dr. Sunando Datta (IISER Bhopal, Bhopal, India). All reagents for immunocytochemistry were purchased through Vector Laboratories Inc (Burlington, ON). RNAscope® Probe targeting Tks5 was purchased through Advanced Cell Diagnostics (Hayward, CA).

### Cell Lines

Human dermal lymphatics endothelial cell line (HDLEC) was purchased from PromoCell (Heidelberg, Germany) and grown in 5% endothelial cell growth medium MV 2. MDA-MB-231 (ATCC HTB-26), MCF-7 (ATCC HTB-22), MDA-MB-231 LNs and HEK293 T cells were grown in DMEM supplemented with 10% fetal bovine serum (FBS; Gibco Life Technologies, Grand Island, NY, USA). 21T-MT cells were cultivated in AMEM (with 10% FBS, 2 mM L-glutamine (Gibco Life Technologies), insulin (1 ug/ml), epidermal growth factor (12.5 ng/ml), hydrocortisone (2.8 mM), 10 mM 4-(2-hydroxyethyl)-1-piperazineethanesulfonic acid (HEPES), 1 mM sodium pyruvate, 0.1 mM non-essential amino acids and 50 ug/mL gentamycin (all the reagents were purchased from Sigma Chemicals). These cells were grown at 37°C and 5% CO2.

For overexpressing Tks5, cells were infected with either lentiviral control or Tks5-GFP particles and were sorted using *BD Influx* cell sorter (BD Biosciences). Cells selection for Tks5-GFP expression was maintained under puromycin (4µg/mL). For Tks5 knockdown, lentivirus containing Tks5 knockdown plasmids were generated. Lentiviral production was performed using HEK293 T cells. Tks5 knockdown plasmids (scrambled control or Tks5-targeting) together with packaging vectors PAX2 and MD2.G were transfected into HEK293 T cells using Lipofectamine 3000 following standard procedures. After 24 and 48 hours, viral particle rich medium was collected, spun down at 300xg to get rid of cellular debris, aliquoted and stored at -80°C. On the day of infection, the cells were treated with viral particles and polybrene (Sigma-Aldrich TR-1003-G). Next day, media was replenished and cells were selected with puromycin (4µg/mL). Tks5 knockdown was confirmed by western blot. For Tks5 knockout, Origene CRISPR/Cas9 Tks5 knockout kit was used. The protocol followed was provided by the manufacturers. CCR7 stable knockdown cell line generation was performed by lentiviral infection and selection as described for that of Tks5 knockdown. CCR7 knockdown was validated by western blot.

### In Vitro Lymphatic Endothelium

An HDLEC monolayer is generated as described previously.^40^ Briefly, HDLEC were cultured (70%) in a fresh medium on a fibronectin (100 µg/mL) coated glass coverslip (No. 1.5, 22 x 22 mm) and supplemented until a monolayer is formed. MDA-MB 231 Tks5-GFP were added on the monolayer and a time-course fixation was performed every 30 minutes for 3 hours. HDLEC were stained, with Zonula occludens 1 (ZO-1) a tight junction marker protein and VE-cadherin, an adherent junction marker protein. Cells were imaged using a Zeiss LSM 700 microscope, with 40x/1.3 Oil DIC numerical aperture objective mounted on a Märzhäuser Scanning Stage D130x100, lymphatics cancer system was monitored for the formation of invadopodia-based protrusions. Images were collected using a high-resolution AxioCam camera and the Zen software program.

For live cell imaging, monolayer was generated in a similar way as described above. Once an intact layer is formed, cells are stained with Cell Mask Orange from Thermofisher (Catalogue number C10045) at 1:10,000 and incubated for 10 minutes. Overnight serum-starved MDA-MB-231 Tks5-GFP stable cells were pre-incubated with Hoechst (1:10,000), for 10 minutes, and added to the monolayer. Cells were imaged every 45 seconds to 2 minutes for up to 2 hours using the high-resolution AxioCam camera and the Zen software program

### In Situ Hybridization and Image Analysis

Breast cancer and matched regional lymph nodes metastatic carcinoma tissue microarray (BR10010f), was obtained from Tissue Arrays (TissueArrays/com, Derwood, MD, USA). The study is compliant with all relevant ethical regulations on the use of human tissue and study approval was obtained from the Institutional Ethics Review Board of UBC (IRB#H17-01442). The TMA was stained for Tks5 mRNA using a Tks5 RNAscope® Probe as previously described.^41^ Briefly, a freshly cut TMA section was baked at 60 °C for 1 h, deparaffinized using Xylene and 100% ethanol, and air-dried. Following this, slides were incubated in retrieval buffer at boiling temperature (100 °C) for 15 min, rinsed in deionized water followed by ethanol, and immediately treated with protease at 40 °C for 30 min (HybEZ-hybridization oven; Advanced Cell Diagnostics, Hayward, CA). Tks5 probe was incubated at 40 °C for 2 h in the HybEZ oven. Slides were washed twice at room temperature and signals were amplified through a 10-step incubation using RNAscope 2.5 HD Reagents Duplex Detection Kit. (ACD, Cat. No. 322435).

Each tissue core was scanned using a Zeiss Axio Observer microscope microscope slide scanner (Carl Zeiss, Germany) at ×20 magnification. Image segmentation and analysis was performed using ZEN Intellesis deep-machine-learning program software (ZEN Intellesis, ZEN 2.6 Blue edition software, Zeiss, Germany). ZEN Intellesis software was used to train and develop a program which could identify nuclei and Tks5 based on pixel color. The program was then applied to each tissue core image and image segmentation was performed. Tumor and stromal areas for each tissue section were manually selected and each area was segmented. Tumor area was used for quantification. The digital output reports the total number of RNA punctate and total number of nuclei. The results are reported as the total counts of RNA punctate divided by the total nuclei counts. This represents the average mRNA expression per cell.

### Immunohistochemistry and Image Analysis

Breast cancer tissue microarray (TissueArray; BR1505E) was obtained and immunostained for CCR7. Briefly, the TMA was baked overnight at 37 °C, followed by deparaffinization in xylene and rehydration using an ethanol gradient followed by a 1Xphosphate-buffered saline (PBS), pH 7.4, wash. For immunostaining heat-induced antigen retrieval was performed at 95 °C for 30 min in a 1mM EDTA with 0.05% tween 20 buffer (pH 8). After heating, slides were cooled to room temperature and washed in 1XPBS. Slides were then incubated with 0.1% Triton X-100 in PBS for 10 min and washed once with 1XPBS for 5min. This was followed by a 30 min incubation in 10% normal goat serum. TMA was washed with PBS and incubated with primary antibody (1:2000 for CCR7, NBP2-67324) at room temperature for 1h. Excess antibody was removed by washing three times with 1XPBS. TMA was incubated with fluorescently labelled secondary antibody (Alexa fluor 647 anti-Rabbit IgG) for 1 h at RT. Excess antibody was removed by washing three times with 1XPBS. To quench autofluorescence, the samples were incubated with 0.1% of Sudan Black for 10min at RT followed by three times washes with PBS. The nuclei were counter stained using Hoechst (1:5000/PBS) for 5min at RT. Excess stain was washed off and the samples were mounted using Prolong Gold mounting medium and left to dry overnight under dark. Images were captured on a Zeiss Axio Observer using a 10X objective and analyzed further using Zeiss Intellesis software.

Each TMA core was scanned using a Zeiss Axio Observer microscope (Carl Zeiss, Germany) at 10X magnification. Image segmentation and analysis was performed using ZEN Intellesis deep-machine-learning program software (ZEN Intellesis, ZEN 2.6 Blue edition software, Zeiss, Germany). ZEN Intellesis software was used to train and develop a program which could identify nuclei and differentiate nuclei and CCR7 staining based on pixel color. The program was then applied to TMA and image segmentation was performed. Tumor and stromal areas for each tissue section were manually selected, and each area was segmented. Tumor area was used for quantification. The digital output reports the total number of nuclei, nuclei area and CCR7 stained area. The results are expressed as CCR7 expression ratio in tumor area (CCR7 stained area (pixel^2^)/nuclei area (pixel^2^).

### Immunoprecipitation

MDA-MB-231LN cells were plated on 10cm plastic dishes which had been coated with unlabeled 0.2% gelatin for 10min at RT and washed with PBS prior to plating. Cells were incubated and allowed to attach for 1h at 37°C, 5%CO2. Following 1h incubation, cells were treated with either 300nM CCL19 or 10µM of CCR7 antagonist (CCR7 Ligand1; 10mM stock made in DMSO) or 1µM of Saracatinib (20mM stock in DMSO) diluted in complete media and incubated for an additional 4h at 37°C, 5%CO2. At the end of the incubation, cells were washed, trypsinized and harvested. Cells were lysed in ice cold RIPA supplemented with 1X HALT Protease and Phosphatase Inhibitor and incubated for 15min at 4°C on a rotamer. The cell lysate was clarified at 12,000xg for 10min at 4°C. The supernatant was collected and kept chilled on ice while quantifying the protein concentration using Pierce Rapid GOLD BCA protein assay kit. For each condition, 1mg of total cell lysate was used for immunoprecipitation which was first precleared at 4°C for 1h using rabbit isotype control and magnetic protein G beads. The precleared lysate was then incubated overnight at 4°C on a rotamer with 1µg of either the target antibody or the isotype control. Following day, 20ul of prewashed magnetic protein G beads were added to the lysate-antibody mixture and incubated for an additional 2h at 4°C on a rotamer. Subsequently, the beads were washed multiple times with ice cold PBS, and the samples were eluted from the beads in SDS sample buffer and immunoblotted for phosphotyrosine.

### GST Pak1-PBD Effector Pull down Assay

GST Pak1-PBD was purified as previously reported. Rac3 activation assay was performed as reported earlier with minor modifications. Briefly, MDA-MB-231 LN cells were plated on 10cm plastic dishes which had been coated with unlabeled 0.2% gelatin for 10min at RT and washed with PBS prior to plating. Cells were allowed to attach for 1h at 37°C, 5% CO2. Following 1h incubation, cells were treated with either 300nM CCL19 or 10µM of CCR7 antagonist (CCR7 Ligand1; 10mM stock made in DMSO) diluted in complete media and incubated for an additional 4h at 37°C, 5%CO2. At the end of the incubation, cells were washed, trypsinized and harvested. Cells were lysed in ice cold lysis buffer (50mM Tris pH 7.5, 10mM MgCl2, 0.5M NaCl, 1% Triton X 100) supplemented with 1X HALT Protease Inhibitor cocktail and 200µM GTP and incubated on ice for 5min. The lysates were clarified at 12,000xg for 10min at 4°C. Cleared lysates (500µg of lysate per condition) were incubated with GST Pak1-PBD (25µg per condition) prebound to magnetic Glutathione Agarose beads for 1.5h at 4°C. Beads were washed three times with ice cold wash buffer (25mM Tris, pH 7.5, 30mM MgCl2, 40mM NaCl). The bound proteins were eluted from the beads using SDS sample buffer and immunoblotted using anti-Rac3 antibody to capture active Rac3 population.

### siRNA Transfection

Cells were seeded 24h prior to siRNA transfection. A standard protocol, available on Thermo Fisher Scientific (https://www.thermofisher.com/ca/en/home/references/protocols/cell-culture/transfection-protocol/stealth-sirna-transfection-lipofectamine) was used using Lipofectamine 3000 as the transfection reagent. MDA-MB-231 LNs were transfected with ON-Target Plus SMART Pool siRNA reagent targeting human VAV2 or Scrambled at 25nM. The sequence for the ON-Target Plus SMART Pool siRNAs against Vav2 are as follows: siRNA #1, 5’-CUGAAAGUCUGCCACGAUA-3’, siRNA #2, 5’-UGGCAGCUGUCUUCAUUAA-3’, siRNA #3, 5’-GUGGGAGGGUCGUCUGGUA-3’, siRNA #4, 5’-GCCGCUGGCUCAUCGAUUG-3’

### Western Blot

Cells were harvested and lysed in ice cold RIPA lysis buffer supplemented with 1X HALT Protease inhibitor cocktail and incubated for 15 min on a rotamer at 4°C. The lysates were clarified at 12,000xg for 10 min at 4°C.Protein concentration was estimated using Rapid GOLD BCA Protein assay kit. Samples were prepared (40 µg of TCL for MDA-MB-231 and 80 µg f TCL MCF-7 for Tks5 blots) by adding 1x SDS sample buffer and boiled at 100°C for 5min. Samples were later separated by SDS-PAGE followed by transfer to nitrocellulose membrane. The membranes were blocked in 5% skim milk for 1h at RT followed by incubation with the respective primary antibodies overnight at 4°C. Next day, membranes were washed with 1X TBS-T and then incubated with biotinylated or fluorescently labeled secondary antibodies diluted in 5% skim milk for 1h and then exposed using Odyssey CLx Imaging System (LI-COR lincoln, NE) or Sapphire FL Biomolecular Imager (Azure Biosystems).

For Phospho antibodies the membranes were blocked using 5% BSA in TBS-T and both primary and secondary antibodies were diluted in 5% BSA in TBS-T.

### Invadopodia Formation Assay/ Gelatin Degradation Assay

The degradation of ECM was observed by coating coverslips with Alexa Fluor 594/568 labeled gelatin as described previously (25,26). Briefly, coverslips were coated with 50 µg/mL of poly L-Lysine for 20 min, followed by treatment with chilled glutaraldehyde (0.5 %) for 15 min. The coverslips were then inverted on a drop of Alexa Fluor 594 or 568 labeled gelatin and incubated for 10min at RT under dark. The coverslips were washed with PBS and then quenched using non-essential amino acids diluted in PBS for 30min at RT. The coverslips were washed with PBS and stored at 4°C under dark for further use. MDA-MB-231 cells (20,000 cells/well) were plated for 6 hours (4 hours for 231-Tks5 GFP). For MDA-MD-231LNs, 1.2x10^5^ cells were plated per well in a 6 well plate on Alexa 568 labelled Gelatin coated coverslip and incubated for 4 or 5h at 37°C. Cells were allowed to attach for 1h prior to addition of CCL19 (300nM) or CCR7 ligand1 (10µM) and further incubated for 2-3h at 37°C, 5%CO2. After incubation, cells were fixed using 4% PFA for 10min followed by washes with PBS. Cells were permeabilized using 0.1% Triton-X-100 for 10min at RT followed by blocking with 5% BSA in PBS. Cells were either stained with primary antibodies (CCR7, 1:500 diluted in blocking solution) for 1hr at RT followed by secondary antibody staining or directly stained for actin using Alexa647 Phalloidin. The coverslips were then mounted on microscopic slides using Prolong® Gold mounting media with DAPI. The slides were dried overnight and imaged using a widefield epifluorescent microscope (Zeiss Axio Observer).

### Migration Assay

Cells were serum starved overnight before being plated (5000 cells/well) in a transwell migration chamber, 8 µm (Costar Corning). Cells were plated diluted in serum free media and fetal bovine serum (20%) was used as a chemoattractant in the bottom chamber. Cells were allowed to migrate 12-16h. Following day, the inserts were washed with PBS and fixed with 4% PFA for 10min at RT. The cells on the top side of the insert were carefully scrapped off using a Q tip and the membranes were sliced out using a blade and mounted on a glass slide using Prolong gold with DAPI. The membranes were imaged using a widefield epifluorescent microscope. The number of cancer cells migrated through the transwell membrane was quantified.

### Lymphatic Invasion Assay

Transwell migration chambers, 8 µm (Costar Corning), were coated with fibronectin (100 µg/mL) at 37°C for 20 minutes. HDLEC cells were cultured on fibronectin layer and supplemented accordingly until an intact monolayer is formed. Cancer cells were serum starved overnight and then added on the intact monolayer (50,000 cells/well). Fetal bovine serum (20%) was added as a chemoattractant in the bottom chamber. The transwells were incubated overnight (36h for 21T MT and 48h for MCF-7). At end-point, the inserts were washed with PBS and fixed with 4% PFA for 10min at RT. The cells on the top side of the insert were carefully scrapped off, membranes excised, and mounted on a glass slide using Prolong gold with DAPI. The membranes were imaged using a widefield epifluorescent microscope. The number of cancer cells on the underside of the transwell membrane was quantified.

### Fibronectin Migration Assay

Transwell migration chambers, 8 µm (Costar Corning), were coated with fibronectin (100 µg/mL) at 37°C for 20 minutes. Cells were serum starved overnight and then added on the fibronectin layer (50, 000 cells/well). Fetal bovine serum (20%) was added as a chemoattractant in the bottom chamber. The cells were allowed to migrate overnight (12-16h). Following day, the inserts were washed with PBS and fixed with 4% PFA for 10min at RT. The cells on the top side of the insert were carefully scrapped off using a Q tip and the membranes were sliced out using a blade and mounted on a glass slide using Prolong gold with DAPI. The membranes were imaged using a widefield epifluorescent microscope. The number of cancer cells migrated through the fibronectin layer alone were quantified.

### Transendothelial Electrical Resistance Assay (TEER)

TEER measurements were made with Epithelial Volt/Ohm (TEER) Meter (World Precision Instruments, Sarasota, FL). Transwell migration chambers, 8 µm were coated with fibronectin (100 µg/mL) and incubated at 37ºC for 20 minutes. HDLEC were cultured in fresh medium on fibronectin-coated-migration chambers until a confluent monolayer was formed. MBA-MB-231 Control, knockout and knockdown cells were added on the layer and TEER measurements were made at various time point (t=0, t=1, t=2 and t=3 hours). Chambers with fibronectin alone were used as negative control. Resistance of HDLEC junctions was calculated as the mean of at least three wells from each experiment (three replicates each) from which the mean resistance of control inserts (fibronectin alone) was subtracted. The final data is presented as percent resistance difference as:

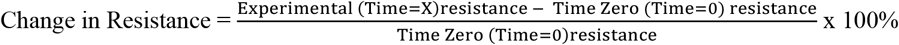

### Inverted Transendothelial Electrical Resistance Assay (TEER)

Invert TEER measurements were made using a similar protocol. Transwell migration chambers, 8 µm were coated with fibronectin (100 µg/mL) and incubated at 37°C for 20 minutes. HDLEC were cultured in fresh medium on fibronectin-coated-migration chambers until a confluent monolayer was formed. The control and mutated MDA-MB-231 cells (100,000/well) were plated at the bottom of the wells for 20 minutes in the form of a drop. After this, the wells were carefully placed back in the same position and the resistance of the trans-endothelial barrier was measured. Here the resistance represents the change in junctional integrity as the cancer cells intravasate through the junctions of the monolayer.

### Proliferation Assay

Cells were seeded in triplicate in 24 well cell culture plates (5000 cells/well) in DMEM + 10% FBS. Immediately after incubation and at 24, 48, and 72 hours the viable cells were measured by measuring their bioluminescence after adding luciferin (150 μg/ml).

### BLI Procedure

MDA-MB-231LN cells stably expressing luciferase were used to generate knockdown and control cells. In vivo BLI was performed using IVIS Lumina scanner (PerkinElmer). Mice were injected intraperitoneally with 100 μL of D-luciferin (30 mg/mL) and were then anesthetized with isofluorane (2% in 100% oxygen) using a nose cone. Whole body BLI imaging (30sec) was used to capture luminescence from the primary tumor. Ex vivo organ imaging was performed to capture luminescence from lymph nodes and lungs. For each image, a standardized region of interest was drawn to encompass the whole tumor or tissue area and the average radiance (photons/sec/cm2/sr) was calculated using Living Image software (PerkinElmer).

### Animal Preparation

All experiments were approved by the Animal Care Committee at The University of British Columbia and were performed in accordance with guidelines established by the Canadian Council on Animal Care. Female NSG mice (aged 6–8 weeks, from Charles River Laboratories, Wilmington, MA) were housed in a pathogen-free modified animal barrier facility. On the day of injection, cells suspended in 0.1 mL of a 1:1 PBS/Matrigel solution were injected into the 2^nd^ mammary fat pad female nude mice, anesthetized with 2% isoflurane in oxygen. The control group (n = 5) and Tks5-KD group (n = 5) were injected with 1million cells per mouse. Sham surgery and lymph node removal surgery was performed on 6 week old female mice. Mice received local anesthetic/analgesic Lidocaine, dose:5 mg/kg at 5 mg/ml and subcutaneous injection at a volume of 20-70ul during surgery and post-surgery. Mice received non-steroidal anti-inflammatory agent Meloxicam (metacam) at a dose of 5 mg/kg by subcutaneous injection post-surgery and each following day for the next two days. 4 days post-surgery, MDA-MB-231LN cells were injected into the 2^nd^ mammary fat pad of the sham surgery mice (n=5) and lymph node removal surgery mice (n=5). All procedures involving animals were approved by the Animal Care Committee at The University of British Columbia and were performed in accordance with guidelines established by the Canadian Council on Animal Care. Animal Care and Ethics Protocol: A21-0237

### Statistical Analysis and Software

Data were handled in Microsoft Excel 16.90.2 and analyzed using GraphPad Prism 10.6.0. Figures were prepared in Adobe Illustrator CC 2025. Data were collected from three biological experimental replicates. Error bars represent the standard error of the mean (SEM). All data sets were assessed for normality using the Kolmogorov–Smirnov or Shapiro Wilk test. Normally distributed data were analyzed using an paired or unpaired t test or Welch’s test. For comparison of two groups or more a one-way ANOVA or two-way ANOVA was performed. P-value was obtained using GraphPad Prism. *p < 0.05. **p < 0.01. ***p < 0.001. ****p < 0.0001.

