## Supplemental Figures for "Invadopodia-Mediated Remodeling of the Lymphatic Endothelium Drives Cancer Cell Lymphatic Dissemination and is Regulated by a CCR7-Vav2-Rac3 Signaling axis"

[*]Equal Contribution

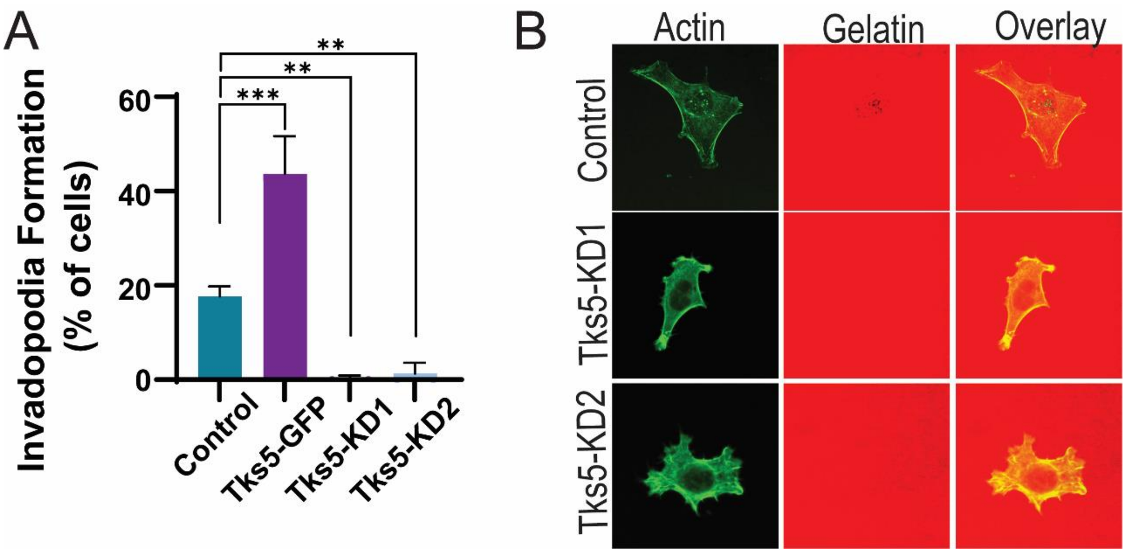


**Supplementary Figure 1: Tks5 regulates invadopodia formation. (A)** Quantification of invadopodia in control, Tks5 overexpressing (Tks5-GFP), and Tks5 knockdown cell lines. Cells were plated on Alexa594-gelatin coated coverslips for 6 h, fixed, permeabilized, and stained with Alexa488-phalloidin to stain Factin. Invadopodium based degradation spots on gelatin, co-localized with F-actin were quantified using confocal microscope. Means ± SEM. N=3. 20 independent spots per sample were counted. Asterix denotes significance measured One-way ANOVA. **(B)** Representative images of invadopodia formation in control cells and loss of formation in Tks5 knockdown cell lines.

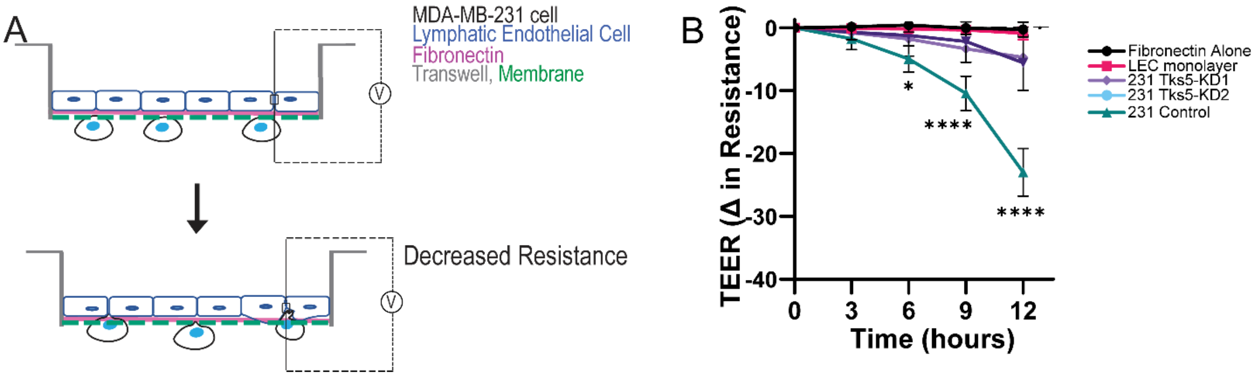


**Supplementary Figure 2: Remodeling of lymphatic endothelial junctions is regulated by Tks5. (A)** Animated diagram depicting cancer cell modification of lymphatic endothelium junctions and the corresponding change to endothelial resistance. **(B)** Transendothelial electrical resistance (TEER) measurements over time for a lymphatic endothelium when incubated with control, Tks5 knockdown cell lines. Two-way Anova. N=3. Mean ± SEM. Asterisk denotes significance.

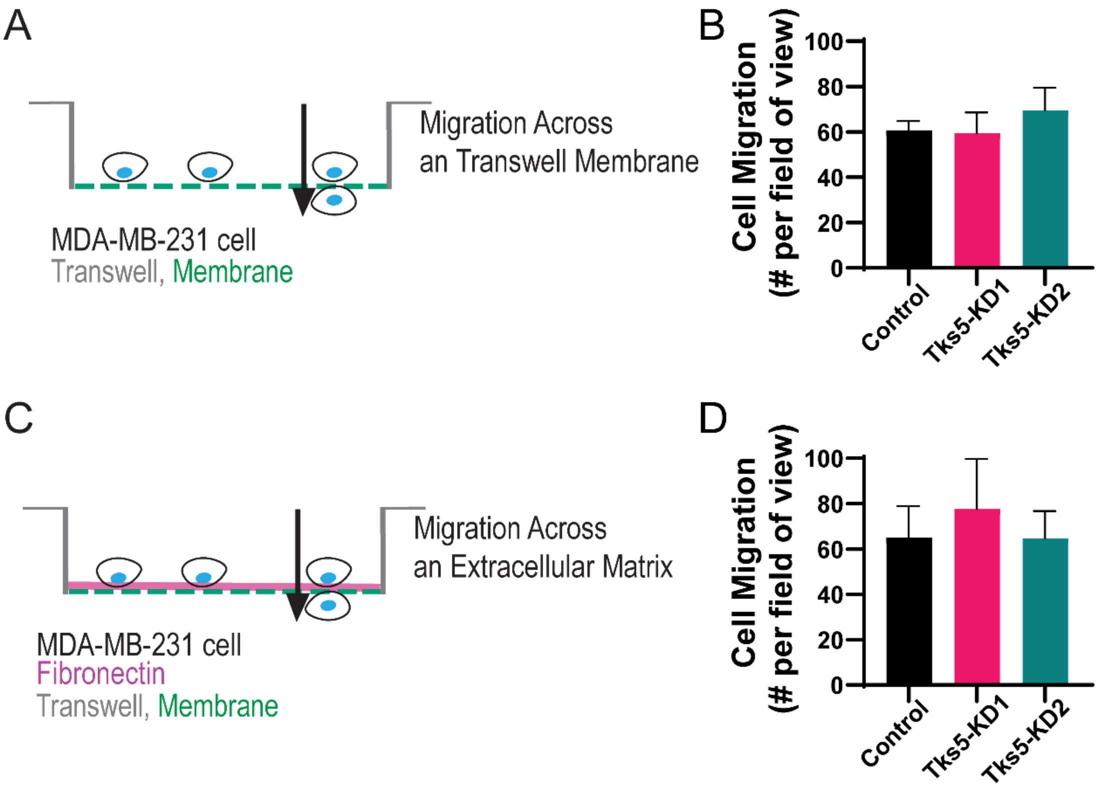


**Supplementary Figure 3: Tks5 does not regulate cell migration. (A)** Schematic representation of cell migration across a transwell. **(B)** Quantification of control and Tks5 knockdown cells able to migrate across a transwell membrane. **(C)** Schematic representation of cells migration across a fibronectin-coated transwell. **(D)** Quantification of control and Tks5 knockdown cells able to migrate across a fibronectin-coated transwell membrane. N=3. ±SEM. Asterix denotes. One-way ANOVA.

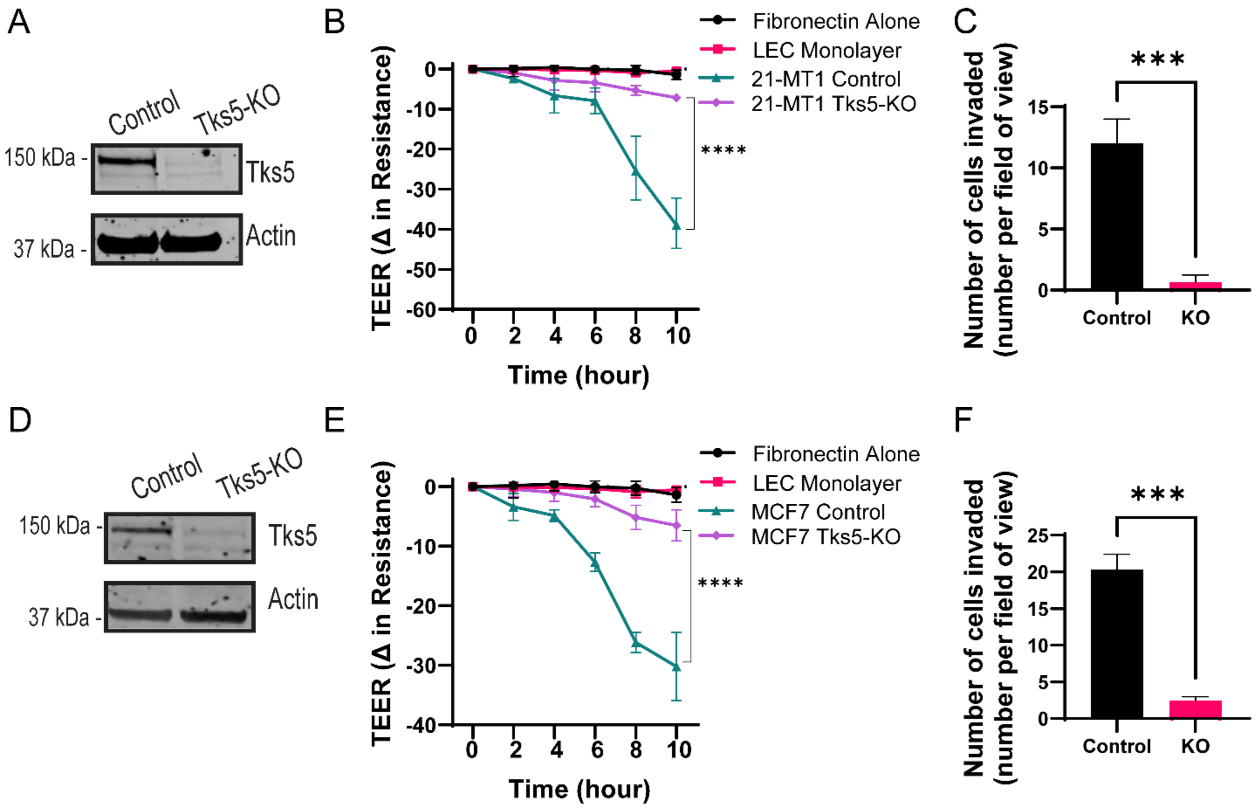


**Supplemental Figure 4: Tks5 regulates breast cancer cell remodeling of lymphatic endothelial junctions and invasion across a lymphatic endothelium.** 21T-MT cells and MCF7 cells were treated with either control or CRISPR Cas9 guide RNA to generate Tks5 knockout cells. Representative western blot of Tks5 levels in (**A**) 21T-MT control and knockout cell lines and **(D)** MCF7 control and knockout cell lines. **(B, E)** Time-dependent transendothelial electrical resistance (TEER) measurements of a lymphatic monolayer cell junctions when incubated with **(B)** 21T-MT control or Tks5 knockout cell lines and **(E)** MCF7 control and Tks5 knockout cell lines for 10 hours. Two-way ANOVA. Invasion of **(C)** 21-MT1 control and Tks5 knockdown cell lines and **(F)** MCF7 control and Tks5 knockout cell lines across a lymphatic endothelium. Student's T-test. N=3. Mean ± SEM. Asterix denotes significance.

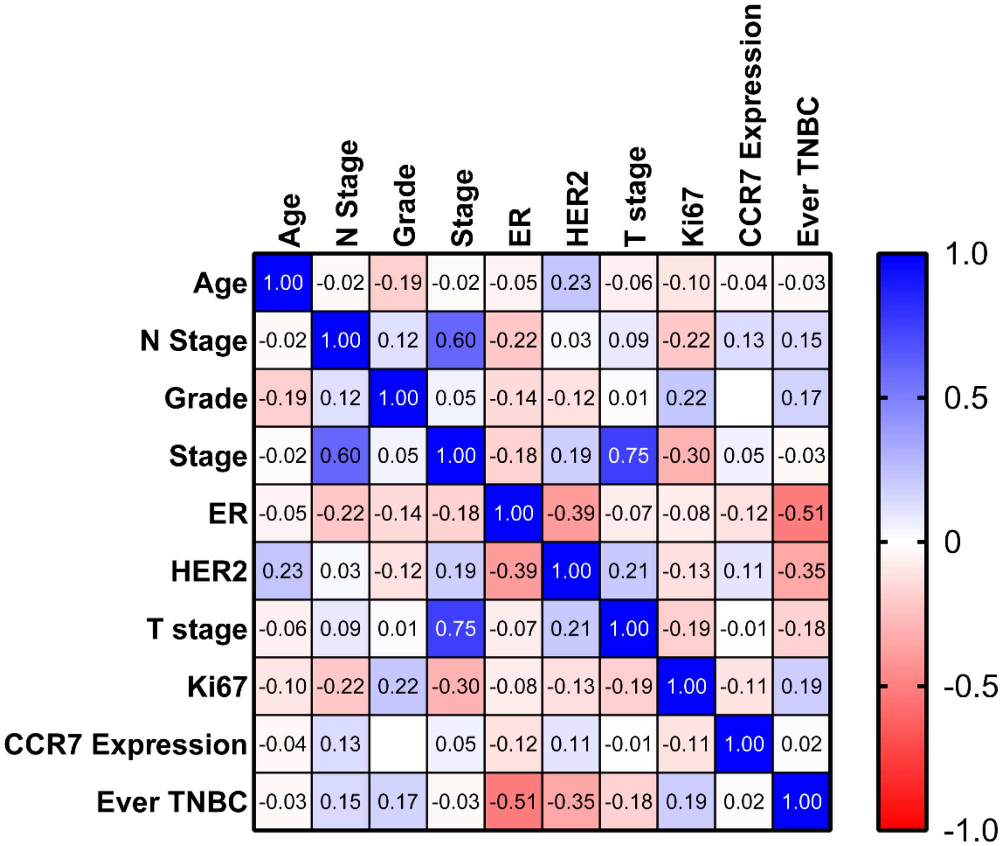


**Supplementary Figure 5. Correlation Map of CCR7 expression and patient clinical data.** Correlation matrix of all clinical data and CCR7 expression. Color map represents correlation strength between each data set; represented as positive (blue) or negative (red). Pearson R values are indicated in each box.

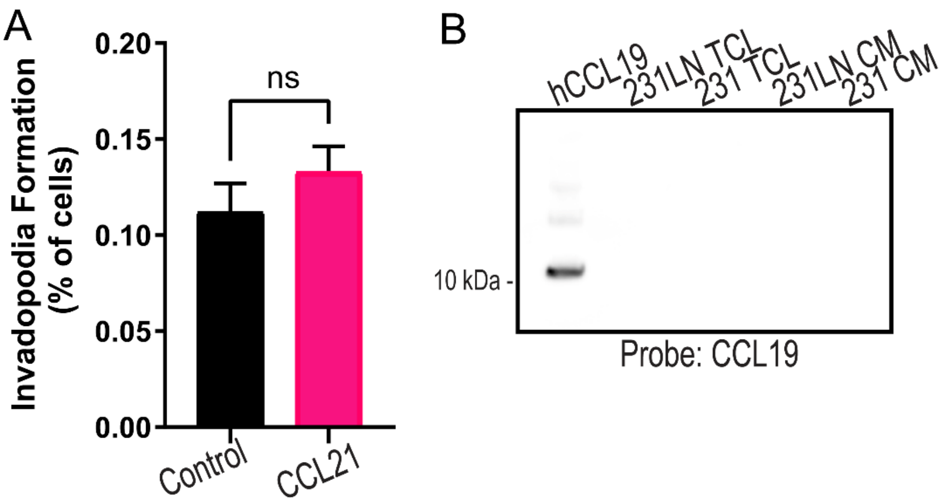


**Supplementary Figure 6. (A)** MDA-MB-231LN invadopodia formation rates in the presence of control (DMSO) or CCL21 ligand. **(B)** CCL19 expression in MDA-MB-231LN and MDA-MB-231 cell lines. Western blot of CCL19 expression in MDA-MB-231LN and MDA-MB-231 cell lines. Human recombinant CCL19 (hCCL19) was used as a control. Tumor cell lysate (TCL) and conditioned media (CM) was analyzed for the presence of CCL19 by western blot.
